# Dynamical Task Switching in Cellular Computers

**DOI:** 10.1101/479998

**Authors:** Angel Goñi-Moreno, Fernando de la Cruz, Alfonso Rodríguez-Patón, Martyn Amos

## Abstract

We present a scheme for implementing a version of task switching in engineered bacteria, based on the manipulation of plasmid copy numbers. Our method allows for the embedding of multiple computations in a cellular population, whilst minimising resource usage inefficiency. We describe the results of computational simulations of our model, and discuss the potential for future work in this area.

## Introduction

Synthetic biology [1,2] is often broadly defined as the *rational engineering* of biological systems, with the aim of implementing novel computational functions in living organisms. Cells such as bacteria may host engineered networks of regulatory proteins - so-called *genetic circuits* - that sense inputs, perform processing, and generate outputs according to human-defined rules.

These artificial cellular computers are often necessarily *single-purpose*, in that they perform one well-defined task, such as the production of drug precursors [3] or the detection of environmental pollutants [4]. Interestingly, the representation of information by physical properties of a biological system (such as levels of gene expression) immediately suggests a parallel with *analogue computers* [5,6], which also used physical quantities (such as hydraulic pressure or the elasticity of a spring) to model specific problems. Importantly, analogue computers were also geared towards specific applications (such as calculating bomb trajectories), and this did not limit their usefulness.

However, we are also interested in the possibility of engineering biological systems that are capable of *task switching* (that is, moving between a number of pre-programmed behaviours (or, “;tasks”) according to specific rules or signals). This allows for the possibility of embedding different tasks into a cellular population, but essentially performing dynamical resource allocation to ensure that the cells do not become metabolically over-burdened. We might compare this to the computer memory management strategy of “paging”, whereby inactive processes are moved from (limited) main memory onto secondary storage (such as a disk). In our model, active processes are represented by plasmids which exist in high numbers, and plasmids that are relatively few in number are considered to be “inactive”.

Here, we present a system in which multiple genetic circuits coexist, and control strategies select which one is functional (i.e., which task *runs*) at a given time. Our method is based on controlling the replication of *plasmids*, which are small DNA molecules which exist (and may be replicated inside) the cell independently of the main chromosomal material.

The rest of this paper is organized as follows: in Section 2 we provide some background to and motivation for our proposed method. In Section 3 we then present our main experimental results, describing our methodology in Section 4. We then conclude in Section 5 with a discussion of our findings, and propose future work.

## Background

Synthetic biology is a rapidly-growing scientific area, with applications in many significant domains, including health, energy, and the environment [7]. One branch of synthetic biology, which we call “cellular computing” [8], is specifically concerned with the construction of computational parts and devices using living cells [9,10]. These implementations include Boolean logic gates [11,12], switches [13], oscillators [14], and counters [15].

A fundamental tool in genetic engineering (and, thus, synthetic biology) are *plasmids*; these are small circular DNA molecules used to introduce new genetic material into bacteria and other cells [16-18]. Typically, new genetic circuits are encoded as a number of genes, the sequences of which are then synthesised and inserted into the plasmid. Plasmids may also naturally be moved *between* cells via *conjugation*, facilitating a process known as *Horizontal Gene Transfer* (HGT) [19,20].

Horizontal gene transfer via conjugation has previously been proposed as a useful mechanism for performing computations [21], by using plasmids to transmit *signals* between cells. Here, we instead embed entire computational circuits within individual plasmids [12], and then manipulate their properties to dynamically switch between them. This allows us to potentially run a number of different “programs” within the cell population, whilst ensuring that only active processes consume scarce system resources. This is the most important aspect of our proposal; while it is, in principle, possible to engineer multiple functional circuits into bacteria (and switch between them), in practice this is difficult to achieve, and inactive circuits place a significant metabolic overhead on the hosts. By dynamically switching between active circuits, and having *only active* circuits present in the host bacteria in significant numbers, we allow for flexible computational behaviour, whilst minimising the burden on the hosts. In the next Section, we describe our model in detail.

### Our task switching model

In previous work, we showed how individual cells may be engineered to exhibit different computational behaviours according to the type of input received [22]. This essentially “flipped” a single genetic circuit between Boolean NAND and NOR, depending on input thresholds. Here, we maintain multiple circuits within a population of cells, and (de)activate them according to need.

A high-level description of our model is shown in Figure 1A. We have two basic levels of control; the lowest level concerns individual genetic circuits encoded in plasmids, and the higher (control) level switches between these circuits (by manipulating the population dynamics of the plasmid pool). We focus mainly on this control level, as the embedding of computational circuits in plasmids is well-understood and standard [12].

**Figure 1.**
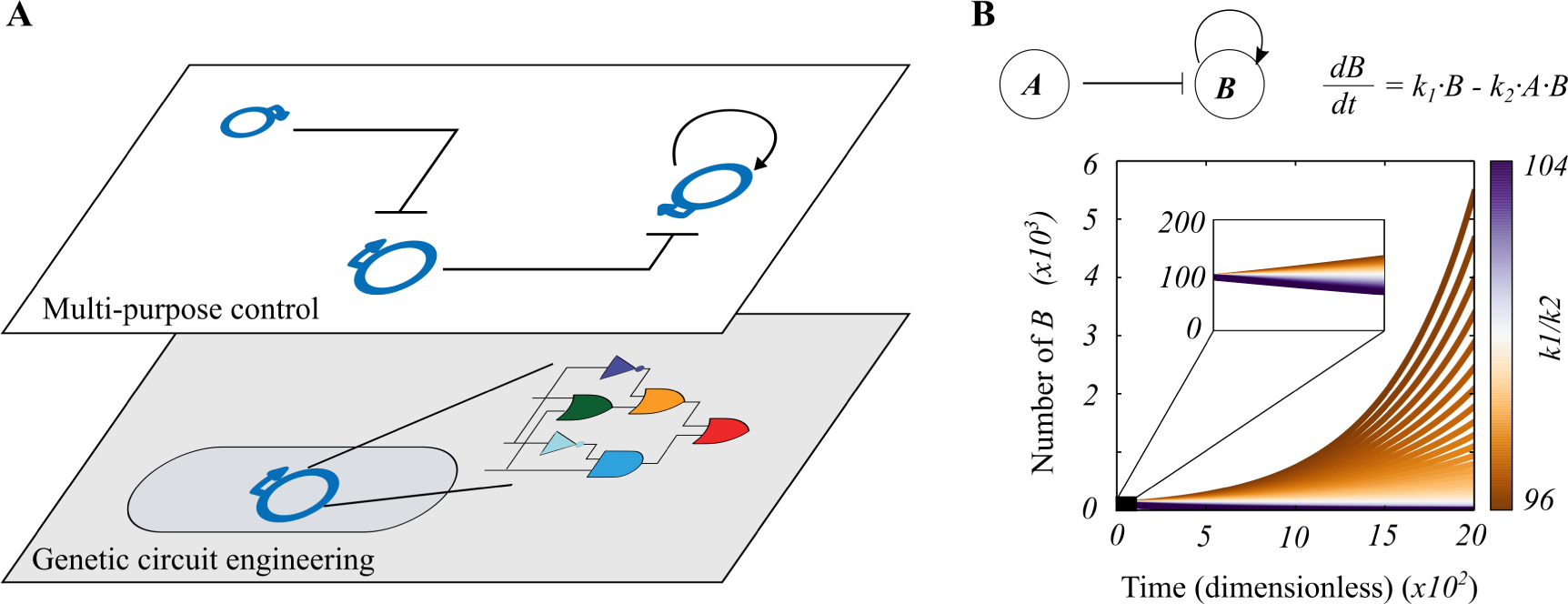
Overall design for a multi-purpose cellular computer. A. The bottom control level encodes a single genetic circuit in a plasmid vector, which executes a single “program”. The top control level handles switching strategies for regulating the numbers of such plasmids in a cellular population. This is done via inhibiting plasmid replication or by promoting plasmid horizontal transfer (i.e. extra replication). B.Deterministic analysis of a two-plasmid system (where plasmid A represses the replication of B, which, in turn, replicates via positive feedback) shows that the system is highly unstable, i.e., plasmid B tends to either increase indefinitely or disappear.

Plasmid replication occurs using the host cell’s DNA replication machinery; plasmids contain sequences known as origins of replication (ori), which “instruct" the host cell to initiate its own replication. Importantly, a plasmid ori sequence may be activated or deactivated by initiator/repressor proteins that bind to it, thus turning on or off the replication of that plasmid [23]. In turn, initiator/repressor proteins may be expressed by genes in other plasmids, allowing one plasmid to effectively turn another on or off. Plasmids propagate in a system through either horizontal transfer via cell-cell conjugation, or vertically, when a cell divides into two daughter cells (Figure 2A. Plasmids that are being repressed are not transmitted vertically, but may still be transferred horizontally between siblings. Two specific attributes of plasmids are of direct interest; their copy number, and their stability. Copy number refers to the expected number of instances of a specific plasmid within a single host cell, and this may be “low” (15-20 copies per cell), “medium” (20-100 copies per cell) or “high” (>500 copies per cell). Engineered plasmids may be “set" to any preselected copy number, but there is an attendant trade-off: the higher the copy number, the higher the metabolic burden on the cell. Plasmid stability [24] exists when, at cell division, each daughter cell receives at least one copy of the plasmid.

**Figure 2.**
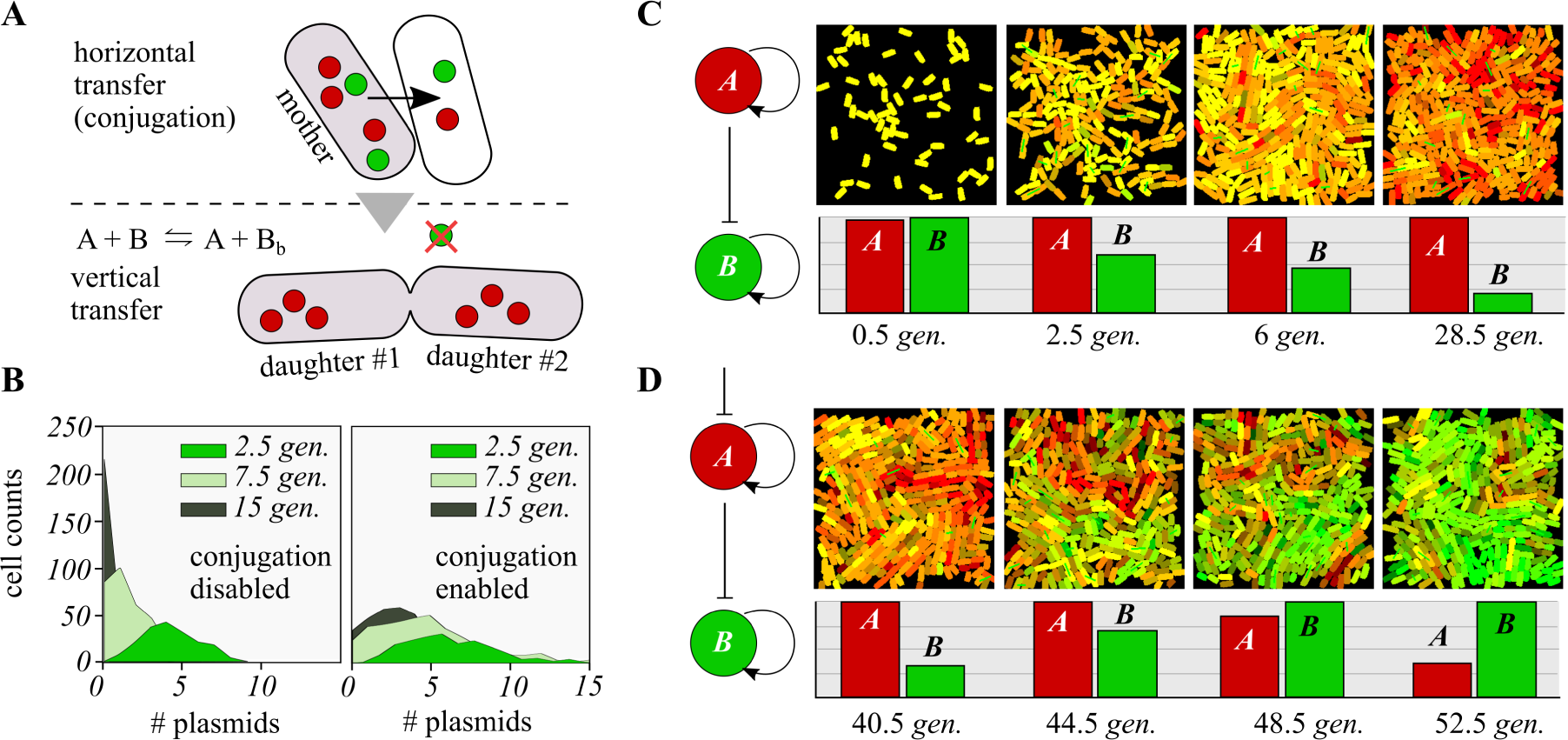
Switchable control of plasmid copy number through horizontal (conjugation) and vertical gene transfer. A. A basic principle guides the following theoretical model: plasmids whose replication is being repressed will not be spread through vertical transfer (i.e. mother to daughters) but can still be copied within siblings via horizontal transfer. B. Distributions representing the number of plasmids in individual cells across a population. The rate of plasmid loss (when its replication is being repressed) is much faster when conjugation is disabled i.e. plasmids are not transferred horizontally. This suggests that conjugation is a powerful tool for stabilizing the systems long enough to allow for switchable computations. C. Simulation of a population where all cells start with two plasmids, A and B. The overall number of B plasmids (bar plots) decreases over time, since its replication is repressed by A. D. Same simulation as in C but the replication of plasmid A can be externally repressed. This repression over A happens after when the overall number of plasmids B is very low (but not zero). As a result, the scenario is reversed and it is plasmid B which predominates over A. Time in all plots is measured in generations(*gen*.) - details on the simulation of conjugation in Methods.

In the next Section, we describe the results of modelling and simulation experiments to investigate the properties and behaviour of systems constructed within our scheme.

## Results

### Continuous modelling

In Figure 1B, we show the behaviour of a two-plasmid system, modelled using ordinary differential equations, in which plasmid A represses (i.e., “turns off”) the replication of plasmid B, which, in turn, replicates via positive feedback. We note a fragile equilibrium for the stability of plasmid B; there is only one scenario, k1/k2 = number of A (100) where the copy number of B does not either increase indefinitely or decrease to zero. This highlights the need for a stabilisation mechanism in such systems, which we describe below.

### Discrete simulation

In Figure 2 we show the results of discrete simulation of a population of cells containing two plasmids, A and B. Each plasmid’s computational “task” is not specified, since we focus here on the dynamics of copy numbers over time (red for plasmid A, and green for plasmid B). We emphasise the fundamental principle that plasmids that are repressed are not spread vertically (through cell division), but may still propagate horizontally (Figure 2A). The significance of this is shown in Figure 2B; for an imagined single plasmid (initial copy number of 10), we consider its representation (in terms of its presence in cells) after a number of periods, with conjugation both disabled (left-hand panel) and enabled (right-hand panel). If conjugation is disabled, then plasmids are essentially rapidly “flushed” from the system, as they are not transferred vertically. However. if we enable conjugation in our simulation, then plasmids are retained for longer within the system, suggesting that conjugation offers an important mechanism for stabilising a system long enough for switchable computations to occur. For details on the simulation of conjugation, please see [25] and the Methods section of the current paper.

In Figure 2C we show the results of spatially-explicit simulations of our system, in which plasmid A represses the replication of plasmid B. Both plasmids start off at roughly equal numbers, but we see that the red plasmid A rapidly dominates the population.

We then show (Figure 2D) how the system may be “switched”, such that an alternative computational task is selected for the population. Replication of plasmid A is repressed by an external signal, which leads to both a gradual loss of the red plasmid A, and an increase in the representation of the green plasmid B (since its replication is no longer being repressed by plasmid A). If we remove the external signal repressing plasmid A, then the system will gradually switch back to a dominant “red” state, and this process is indefinitely repeatable.

The spatially-explicit nature of our simulation means that it is possible to analyse further the distribution of plasmids in the system. In Figure 3 we show a snapshot of a simulation in which plasmid B (green) predominates. We see that the distribution of plasmid B is certainly not uniform, and observe clusters that are suggestive of both vertical and horizontal transfer. This clustering is responsible for stabilising the simulation, compared to the situation shown in Figure 1B (i.e., parameter values do not need to be unique). Clusters essentially act like plasmid reservoirs and generate non-linear dynamics around overall plasmid copy number, which favours robustness. In addition, these clusters add a different viewpoint to the analysis of bacterial differentiation within a population [26] - often a knowledge gap - which is advantageous to us.

**Figure 3.**
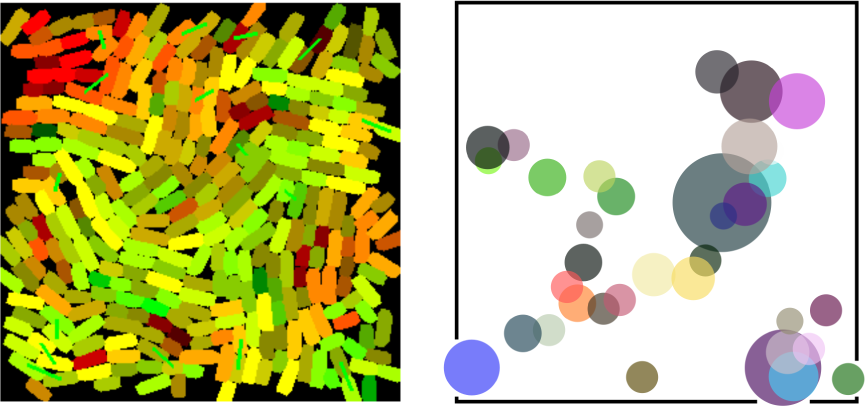
Plasmid copy numbers are not homogeneously distributed across a population,but highly clustered. Dots in the right-hand plot show the positions of those cells with more of plasmid B (green) in the simulation snapshot shown to the left. There is one dot per cell with high B concentration; the colour of dots is meaningless; diameter of dots are directly proportional to the plasmid copy number in each cell. Some dots are perfectly aligned, which suggest vertical transfer, while groups of cells (for example, in the bottom right) increased the copy number via conjugation.

### Distributed computations using cell consortia

An important recent development in synthetic biology and biocomputing has been the development of computational consortia; that is, computations that are distributed over a number of different cells, each of which performs a specific role [27-29]. This approach potentially allows for much more scalable cellular computation, as a large and potentially complex circuit may be broken down into smaller communicating components, each of which is placed in a specific cell. A common structure involves “sender” and “receiver” bacterial strains, each of which either transmit or act upon specific signals, and we adopt that model here.

In Figure 4 we depict our scheme for multi-cellular computation of the Boolean NOR function, using four cell strains that interact to evaluate the gate. Recall that NOR is a negated OR function, so it returns “1” only when both of its inputs are zero, and “0” in all other cases (usefully, NOR is a universal gate, which means that any other Boolean function may be constructed using it).

**Figure 4.**
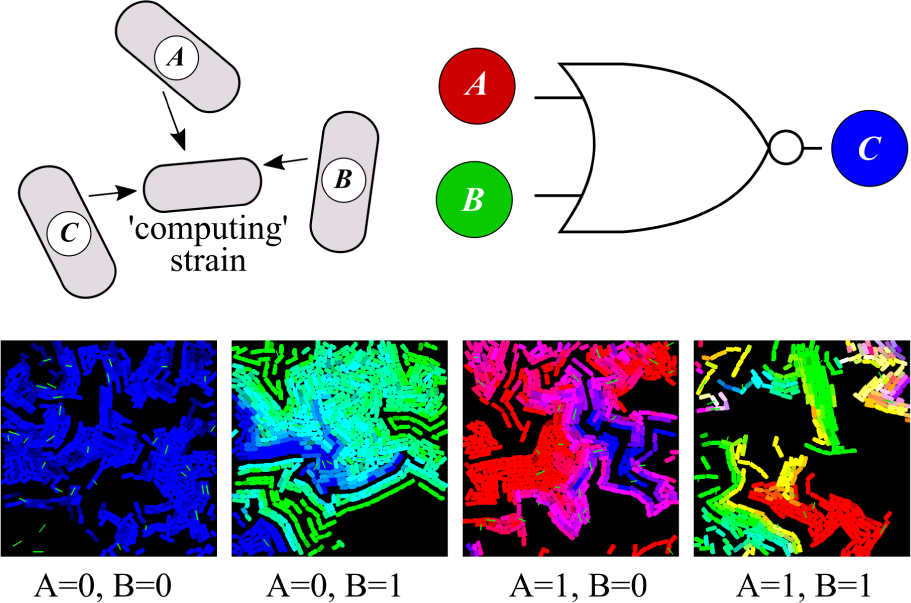
Multicellular computation in a 4-strain consortia. A different approach is adopted to design a 3-plasmid system that responds to a NOR logic function: both input plasmids A and B repress the replication of output plasmid C. We have one strain per plasmid, plus another computing strain - the input/output strains are able to transfer plasmids horizontally to the computing strain but cannot receive plasmids from others. Simulations of the four logic cases highlight the spatial localization of the computation. Only the computing strain is shown - black spaces in-between correspond to different “sender" strains.

Our system is composed of three plasmids; A (red) and B (green) represent the inputs to the NOR gate, and plasmid C (which defaults to blue) representing output=1. Both A and B repress the output plasmid C, so C is only present (corresponding to an output value of “1”) if both A and B are absent (i.e., both input values are equal to zero). Each plasmid is represented by its own bacterial strain (the input plasmids in the “sender” strains), and we also use a fourth “computing” (or “receiver”) strain, which is engineered to express the appropriate fluorescent protein, according to the plasmid that it receives.

We show the results of simulations for each of the four input cases (00, 01, 10 and 11); in the first case, we see only blue cells, as that is the only situation in which we can expect to see an output value of 1. In the other cases, we see a preponderance of green (where the B input dominates), red (where the A input dominates), or a mixture (where both input plasmids are represented equally). This confirms the in-principle possibility of engineering plasmid copy numbers for the purposes of distributed cellular computation.

### Usecasing the potential of task switching

We developed two simple models to demonstrate the potential of the suggested strategy (Figure 5). There are two different approaches to the use of task switching: (1) switch between two completely different tasks, and (2) repurpose the meaning of the inputs to the same task.

**Figure 5.**
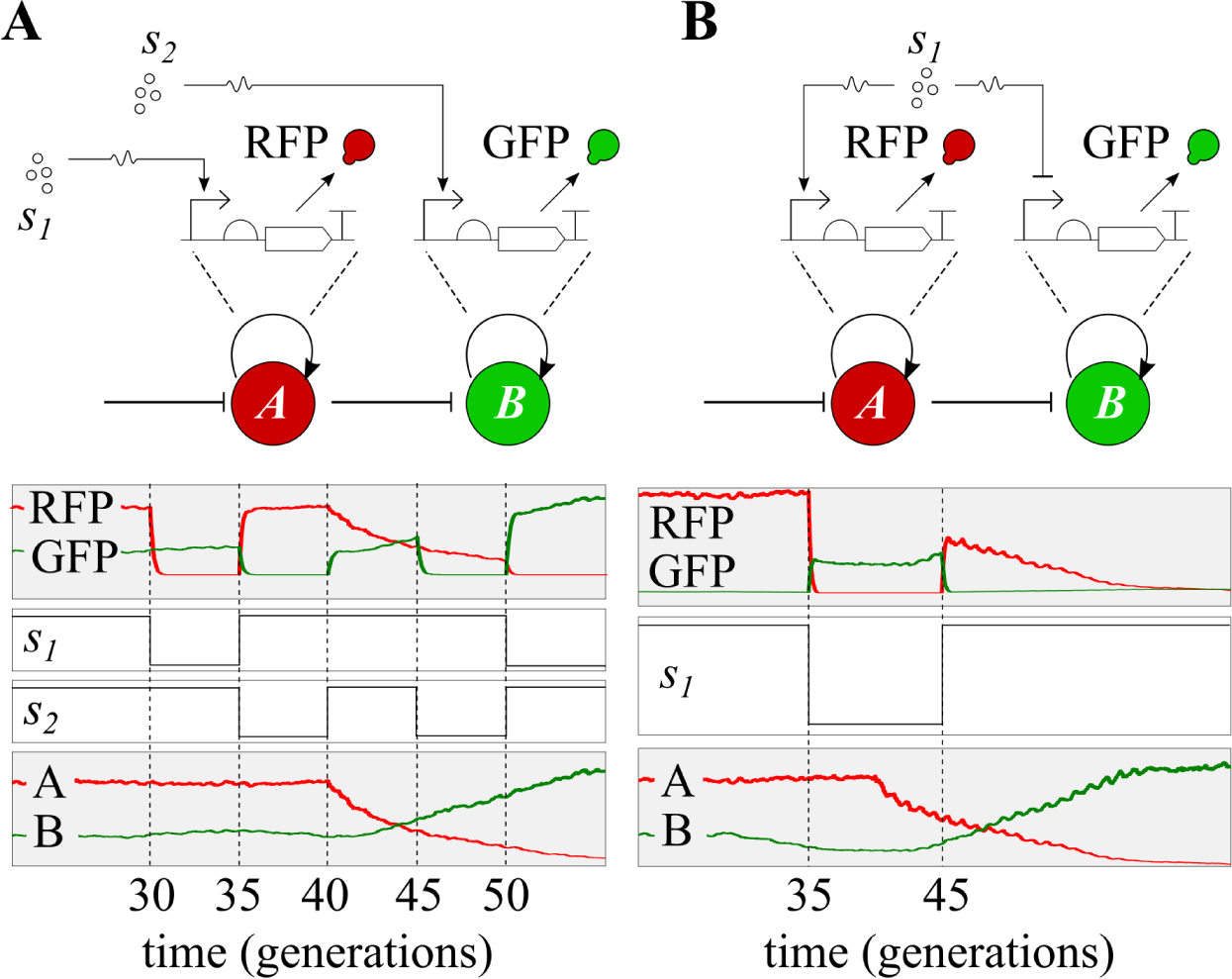
Usecasing the potential of multitasking. A. Reduction of cellular workload. Two input signals, s1 and s2, trigger the expression of different circuits - in this case a simple reporter gene. By controlling plasmid copy number, the population reacts to one of either two input signals. Cells will not have both circuits at the same time, thus reducing the metabolic cost. B. Repurposing input signals. One input, s1, acts as an inducer for the circuit in plasmid A and a repressor for the circuit in plasmid B. Multitasking control allows for switching the population from using s1 as an inducer to using s2 as an inhibitor (and vice-versa).

In the population simulated in Figure 5A, two plasmids coexist, each encoding an different inducible promoter with a fluorescent reporter downstream (red for plasmid A, green for plasmid B). These two tasks are different in that they respond to different input signals (s1 and s2 respectively). Depending on which task runs at a given time, the cells will be sensing the corresponding signal. Therefore, this approach allows us to encode different tasks (e.g., biosensors) that will be active on demand. Figure 5A shows the performance of the simulation over 50 generations. Plasmids A are predominant until t =40, when they are externally repressed; as a result, plasmids B take their place. Simultaneously, the two input signals are changed over time; note that values 0 and 1 indicate their absolute absence and their saturation, respectively. We see that during the first 40 generations, only the dynamics of signal s1 are captured by the population, while signal s2 is ignored or captured at residual levels. The reverse situation occurs from t = 40 onwards, when only signal s2 is sensed.

The repurposing of the device (i.e., the same implementation, but different functions) is illustrated in Figure 5B, by changing the effects of the input on the circuit. Similar tasks as before coexist in a population, where both plasmids express a reporter. In this example, there is only one input signal, which is an inducer for the circuit in plasmid A and a repressor for that of plasmid B. By changing the meaning of inputs, the computation returns a different output. The simulated population reads the signal as an inducer until t = 40, when plasmid A is externally repressed. From then on, plasmids B will be present in higher copy numbers, and the same signal will be read as a repressor.

## Materials and Methods

### Differential models

Ordinary Differential Equations (ODEs) were used to perform deterministic simulations of plasmid dynamics in the non-spatial (Figure 1) and spatial (Figure 5) scenarios. The first set of ODEs describe the reactions 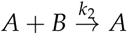 and 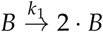

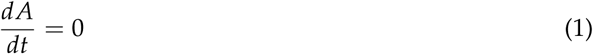

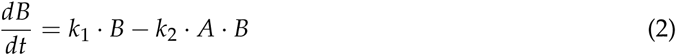

where plasmid A is constant, k_1_ = 0.05, k_2_ = 0.005, and initial conditions are A = B = 100 (all units dimensionless). Equilibrium is found at k_1_/k_2_ = 100 (Figure 1).

The circuits of Figure 5 were also simulated deterministically. Both models run inside cells of a spatial, discrete simulation. The set of ODEs that govern the performance of the first circuit (Figure 5A) is:

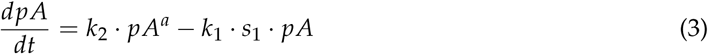

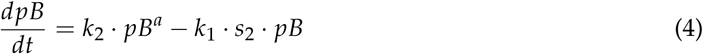

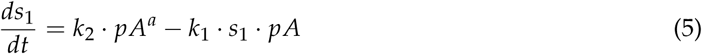

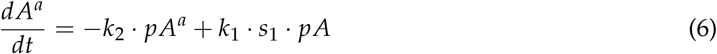

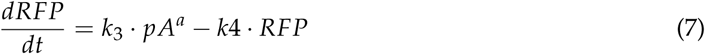

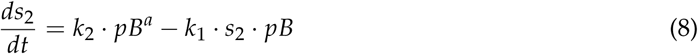

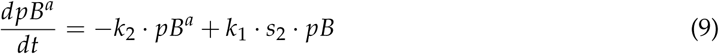

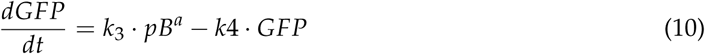

where *pA* and *pB* are the promoter in plasmids A and B, respectively, *k*_1_ = 1 is the rate of binding of the signals to their cognate promoters in either plasmid, denoted by *pA^b^* or *pB^b^, k_2_* = 50 rates the reversed reaction (unbinding) back to pA or pB, k_3_ = 200 is the expression (merged transcription and translation) of the target gene in each plasmid, and k_4_ = 1 the degradation of the proteins GFP and RFP. The number of plasmids A and B is determined by the discrete simulation.

The circuit in Figure 5B was based on the same equations as above, but with the following changes: signal *s2* is removed from the system, signal *s1* inhibits *B* and GFP is expressed from *pB* rather than *pB^a^*. Therefore, equations 5 and 10 change into:

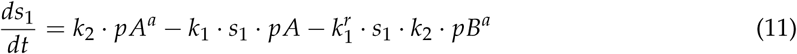

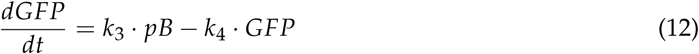

where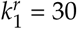, which is the rate of repression of pB by signal s_1_. All rates are expressed in molecules and hours, following values commonly used in mathematical models [30,31]. In any case, signals s1 and s2 are abundant or absent, therefore their derivatives can be considered null.

### Stochastic models

Gillespie’s algorithm [32] was used to calculate the intracellular performance of plasmid stability in Figures 2-4. In Figures 2-3 the reactions simulated were 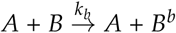 and its reversed 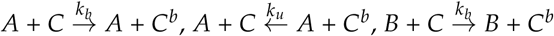, where k_b_ is the rate of plasmid A to block the replication of B, and k_u_ is the rate of unblocking such repression. Their values are 1 and 0.5 respectively.

Figure 4 includes a third plasmid, C to perform a NOR logic function. The reactions for this simulation are: 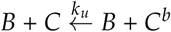

### Spatial simulations

For spatially-explicit simulations we used the agent-based tool DiSCUS [25]. This platform has previously been used to study the spread and growth of bacterial populations [33], and has been included in design-build-test synthetic biology life cycles [34]. In DiSCUS, a population of rod-shaped cells grows on a 2D surface. Each cell was coded to run a copy of either the stochastic or deterministic simulation under study. The spatial simulation resolved plasmid loss due to vertical transfer and plasmid gain due to horizontal transfer. After each division and conjugation event, plasmid copy numbers were updated, and the intracellular simulations adjusted the final numbers of repressed elements accordingly.

Time was measured in theoretical doubling-times, what we called generations (Figure 2), which is the time it takes for a rod-shaped body to grow and divide in DiSCUS. During each conjugation event, 2 or 3 (random) plasmids are transferred. Conjugation frequencies where fitted to experimental observations (see [25] for details). DiSCUS implements the probability that a cell will conjugate with a neighbour cell at any time-point during its lifetime (probabilities range from 0.001 to 0.05). This parameter was fitted to frequencies obtained experimentally, both in liquid cultures [35] and on 2D surfaces [36].

## Discussion and conclusions

The engineering of increasingly complex tasks (i.e., genetic circuits) in cells is a major challenge, and a very active research topic. However, the design of management strategies for the execution of these tasks has received relatively little attention. Computer science, commonly used to frame the development of genetic circuits, has successfully achieved strategies to this end that can be of use to synthetic biology. Here, we present a task switching method in bacteria as a way of managing cellular resources.

Task switching is designed by controlling plasmid copy number (CN). Since each plasmid will encode for a specific task, the control of CN will result in the population running one task or another, without the need to re-design and re-engineer the cells. As envisioned here, this control is achieved via transcription factors [23]; that is, by making plasmid replication dependent on a repressor (for instance, using the LacI repressor to inhibit the replication of a replicon modified with the LacI operator (LacO)). The control of the repressor-operator interplay (e.g. fine-tuning the level of repressor or the noise patterns [37]) can avoid a situation that ends in plasmid loss. We use Horizontal Gene Transfer (HGT) as a tool to solve this issue. According to our simulations, HGT generates plasmid reservoirs that restore the equilibrium to an otherwise collapsing (i.e., inevitable plasmid loss) scenario. Another possibility would be to engineer a second replicon, corresponding to low CN, on the plasmids. This way, plasmids will never be completely lost. Nevertheless, the potential control mechanisms over HGT [38] means that it lends itself to both single-strain and multicellular computations based on this approach.

Plasmids and HGT may play a major role in interbacterial relationships and the evolution of microbial communities [39]. Such powerful tools should not be left out of the synthetic biology toolbox. This study demonstrates the in-principle feasibility of using them to achieve complex human-defined computations in cellular systems, and provides baseline information for their future wet-lab implementation.

## Acknowledgements

The work of AG-M is supported by the SynBio3D (UK-EPSRC-EP/R019002/1) project of the UK Engineering and Physical Sciences Research Council and the BioRoboost (EU-H2020-BIOTEC-820699) Contract of the European Union. The work of AR-P is supported by the Spanish TIN2016-81079-R, (MINECO AEI/FEDER, EU) and Madrid Gov. project B2017/BMD-3691, InGEMICS-CM (FSE/FEDER, EU). Work in FdlC laboratory was financed by grant BFU2017-86378-P from the Ministry of Science and Technology (Spain).

## Author contributions

AG-M, FdlC, AR-P and MA conceived the study and contributed to paper writing. AG-M set up the computational framework and performed all simulations.

## References

1. Amos, M.; Goni-Moreno, A. Cellular computing and synthetic biology. In Computational Matter; Stepney, S.; Rasmussen, S.; Amos, M., Eds.; Springer, 2018; pp. 93–110.

2. Church, G.M.; Elowitz, M.B.; Smolke, C.D.; Voigt, C.A.; Weiss, R. Realizing the potential of synthetic biology. Nature Reviews Molecular Cell Biology 2014, 15, 289.

3. Ro, D.K.; Paradise, E.M.; Ouellet, M.; Fisher, K.J.; Newman, K.L.; Ndungu, J.M.; Ho, K.A.; Eachus, R.A.; Ham, T.S.; Kirby, J.; others. Production of the antimalarial drug precursor artemisinic acid in engineered yeast. Nature 2006, 440, 940.

4. Kim, H.J.; Jeong, H.; Lee, S.J. Synthetic biology for microbial heavy metal biosensors. Analytical and Bioanalytical Chemistry 2018, pp. 1–13.

5. Selvester, R.H.; Collier, C.R.; Pearson, R.B. Analog computer model of the vectorcardiogram. Circulation 1965, 31, 45–53.

6. Heinmets, F. Analog computer analysis of a model-system for the induced enzyme synthesis. Journal of Theoretical Biology 1964, 6, 60–75.

7. Wintle, B.C.; Boehm, C.R.; Rhodes, C.; Molloy, J.C.; Millett, P.; Adam, L.; Breitling, R.; Carlson, R.; Casagrande, R.; Dando, M.; others. Point of View: A transatlantic perspective on 20 emerging issues in biological engineering. Elife 2017, 6, e30247.

8. Amos, M., Ed. Cellular Computing; Oxford University Press, 2004.

9. Manzoni, R.; Urrios, A.; Velazquez-Garcia, S.; de Nadal, E.; Posas, F. Synthetic biology: insights into biological computation. Integrative Biology 2016, 8, 518–532.

10. Benenson, Y. Biomolecular computing systems: principles, progress and potential. Nat. Rev. Genet. 2012, 13, 455–468.

11. Bonnet, J.; Yin, P.; Ortiz, M.E.; Subsoontorn, P.; Endy, D. Amplifying genetic logic gates. Science 2013, 340, 599–603.

12. Nielsen, A.A.; Der, B.S.; Shin, J.; Vaidyanathan, P.; Paralanov, V.; Strychalski, E.A.; Ross, D.; Densmore, D.; Voigt, C.A. Genetic circuit design automation. Science 2016, 352, aac7341.

13. Lou, C.; Liu, X.; Ni, M.; Huang, Y.; Huang, Q.; Huang, L.; Jiang, L.; Lu, D.; Wang, M.; Liu, C.; Chen, D.; Chen, C.; Chen, X.; Yang, L.; Ma, H.; Chen, J.; Ouyang, Q. Synthesizing a novel genetic sequential logic circuit: a push-on push-off switch. Molecular Systems Biology 2010, 6.

14. Purcell, O.; Savery, N.J.; Grierson, C.S.; di Bernardo, M. A comparative analysis of synthetic genetic oscillators. J. R. Soc. Interface 2010, 7, 1503–24.

15. Friedland, A.E.; Lu, T.K.; Wang, X.; Shi, D.; Church, G.; Collins, J.J. Synthetic gene networks that count. Science 2009, 324, 1199–1202.

16. Funnell, B.E.; Phillips, G.J. Plasmid Biology; Vol. 672, ASM Press, 2004.

17. Smillie, C.; Garcillán-Barcia, M.P.; Francia, M.V.; Rocha, E.P.; de la Cruz, F. Mobility of plasmids. Microbiology and Molecular Biology Reviews 2010, 74, 434–452.

18. Martínez-García, E.; Aparicio, T.; Goñi-Moreno, A.; Fraile, S.; de Lorenzo, V. SEVA 2.0: an update of the Standard European Vector Architecture for de-/re-construction of bacterial functionalities. Nucleic Acids Research 2014, 43, D1183–D1189.

19. Llosa, M.; Gomis-Rüth, F.X.; Coll, M.; Cruz, F.d.l. Bacterial conjugation: a two-step mechanism for DNA transport. Molecular Microbiology 2002, 45, 1–8.

20. Tatum, E.; Lederberg, J. Gene recombination in the bacterium *Escherichia coli*. Journal of Bacteriology 1947, 53, 673.

21. Goñi-Moreno, A.; Amos, M.; de la Cruz, F. Multicellular computing using conjugation for wiring. PLoS ONE 2013, 8, e65986.

22. Goñi-Moreno, A.; Amos, M. A reconfigurable NAND/NOR genetic logic gate. BMC Systems Biology 2012, 6, 126.

23. Gil, D.; Bouché, J.P. ColE1-type vectors with fully repressible replication. Gene 1991, 105, 17–22.

24. Friehs, K. Plasmid copy number and plasmid stability. In New Trends and Developments in Biochemical Engineering; Scheper, T., Ed.; Springer Berlin Heidelberg, 2004; Vol. 86, Advances in Biochemical Engineering, pp. 47–82.

25. Goni-Moreno, A.; Amos, M. DiSCUS: a simulation platform for conjugation computing. International Conference on Unconventional Computation and Natural Computation. Springer, 2015, pp. 181–191.

26. García-Betancur, J.C.; Goñi-Moreno, A.; Horger, T.; Schott, M.; Sharan, M.; Eikmeier, J.; Wohlmuth, B.; Zernecke, A.; Ohlsen, K.; Kuttler, C.; others. Cell differentiation defines acute and chronic infection cell types in *Staphylococcus aureus*. Elife 2017, 6, e28023.

27. Goñi-Moreno, A.; Redondo-Nieto, M.; Arroyo, F.; Castellanos, J. Biocircuit design through engineering bacterial logic gates. Natural Computing 2011, 10, 119–127.

28. Regot, S.; Macia, J.; Conde, N.; Furukawa, K.; Kjellén, J.; Peeters, T.; Hohmann, S.; de Nadal, E.; Posas, F.; Solé, R. Distributed biological computation with multicellular engineered networks. Nature 2011, 469, 207.

29. Macía, J.; Posas, F.; Solé, R.V. Distributed computation: the new wave of synthetic biology devices. Trends in Biotechnology 2012, 30, 342–349.

30. Miró-Bueno, J.M.; Rodríguez-Patón, A. A simple negative interaction in the positive transcriptional feedback of a single gene is sufficient to produce reliable oscillations. PloS One 2011, 6, e27414.

31. Goni-Moreno, A.; Amos, M. Model for a population-based microbial oscillator. BioSystems 2011, 105, 286–294.

32. Gillespie, D.T. Exact stochastic simulation of coupled chemical reactions. The Journal of Physical Chemistry 1977, 81, 2340–2361.

33. Espeso, D.R.; Martínez-García, E.; de Lorenzo, V.; Goñi-Moreno, Á. Physical forces shape group identity of swimming *Pseudomonas putida* cells. Frontiers in Microbiology 2016, 7, 1437.

34. Goñi-Moreno, A.; Carcajona, M.; Kim, J.; Martínez-García, E.; Amos, M.; de Lorenzo, V. An implementation-focused bio/algorithmic workflow for synthetic biology. ACS Synthetic Biology 2016, 5, 1127–1135.

35. del Campo, I.; Ruiz, R.; Cuevas, A.; Revilla, C.; Vielva, L.; de la Cruz, F. Determination of conjugation rates on solid surfaces. Plasmid 2012, 67, 174–182.

36. Seoane, J.; Yankelevich, T.; Dechesne, A.; Merkey, B.; Sternberg, C.; Smets, B.F. An individual-based approach to explain plasmid invasion in bacterial populations. FEMS Microbiology Ecology 2011, 75, 17–27.

37. Goñi-Moreno, A.; Benedetti, I.; Kim, J.; de Lorenzo, V. Deconvolution of gene expression noise into spatial dynamics of transcription factor–promoter interplay. ACS Synthetic Biology 2017, 6, 1359–1369.

38. Zatyka, M.; Thomas, C.M. Control of genes for conjugative transfer of plasmids and other mobile elements. FEMS Microbiology Reviews 1998, 21, 291–319.

39. van de Guchte, M. Horizontal gene transfer and ecosystem function dynamics. Trends in Microbiology 2017, 25, 699–700.

